# Metabolic profiling of alcohol consumption in 9778 young adults

**DOI:** 10.1101/041582

**Authors:** Peter Würtz, Sarah Cook, Qin Wang, Mika Tiainen, Tuulia Tynkkynen, Antti J. Kangas, Pasi Soininen, Jaana Laitinen, Jorma Viikari, Mika Kähönen, Terho Lehtimäki, Markus Perola, Stefan Blankenberg, Tanja Zeller, Satu Männistö, Veikko Salomaa, Marjo-Riitta Järvelin, Olli T. Raitakari, Mika Ala-Korpela, David A Leon

## Abstract

**Background:** High alcohol consumption is a major cause of morbidity, yet alcohol is associated with both favourable and adverse effects on cardiometabolic risk markers. We aimed to characterize the associations of usual alcohol consumption with a comprehensive systemic metabolite profile in young adults.

**Methods:** Cross-sectional associations of alcohol intake with 86 metabolic measures were assessed for 9778 individuals from three population-based cohorts from Finland (age 24-45 years, 52% women). Metabolic changes associated with change in alcohol intake during 6-year follow-up were further examined for 1466 individuals. Alcohol intake was assessed by questionnaires. Circulating lipids, fatty acids and metabolites were quantified by high-throughput nuclear magnetic resonance metabolomics and biochemical assays.

**Results:** Increased alcohol intake was associated with cardiometabolic risk markers across multiple metabolic pathways, including higher lipid concentrations in HDL subclasses and smaller LDL particle size, increased proportions of monounsaturated fatty acids and decreased proportion of omega-6 fatty acids, lower concentrations of glutamine and citrate (P<0.001 for 56 metabolic measures). Many metabolic biomarkers displayed U-shaped associations with alcohol consumption. Results were coherent for men and women, consistent across the three cohorts, and similar if adjusting for body mass index, smoking and physical activity. The metabolic changes accompanying change in alcohol intake during follow-up resembled the cross-sectional association pattern (*R*^2^=0.83, slope=0.72±0.04).

**Conclusions:** Alcohol consumption is associated with a complex metabolic signature, including aberrations in multiple biomarkers for elevated cardiometabolic risk. The metabolic signature tracks with long-term changes in alcohol consumption. These results elucidate the double-edged effects of alcohol on cardiovascular risk.

**Key messages:**

- Alcohol intake in young adults associates with multiple novel metabolic biomarkers for the risk of cardiovascular disease and type 2 diabetes. The metabolic aberrations are mainly adversely related to cardiometabolic risk
- Prominent metabolic associations with alcohol consumption include monounsaturated fatty acids, omega-6 fatty acids, glutamine, citrate and lipoprotein particle size. Many of these cardiometabolic biomarkers were as strongly associated with alcohol intake as HDL cholesterol
- The strongest novel biomarkers of alcohol consumption followed linear association shapes, whereas many lipid measures displayed U-shaped associations
- The detailed metabolic signature of alcohol consumption tracked with long-term changes in alcohol use, suggesting that the observed metabolic changes are at least partly due to alcohol consumption
- The results provide improved understanding of the diverse molecular processes related to alcohol intake. Novel metabolic biomarkers reflecting both alcohol intake and cardiovascular risk could serve as intermediates that may help to bridge the complex relation between alcohol and cardiometabolic risk

## INTRODUCTION

Alcohol consumption is one of the leading risk factors for death and disability, accounting for almost 3 million annual deaths globally and 3.9% of life years lost to disease.^1^ The harmful effects of alcohol use are well characterized and uncontentious for some conditions, such as liver cirrhosis, injuries, and several cancers.^1,2^ However, considerable debate remains as to the nature of the association between the amount of alcohol consumed and cardiovascular disease.^3-5^ Observational studies have generally found a U-shaped relationship between alcohol intake and cardiovascular disease risk, with lower risk in moderate drinkers compared to both abstainers and heavier drinkers.^3^ Whether this effect is causal or due to confounding remains an unresolved issue. A recent Mendelian randomization study suggested adverse effects of any amount of alcohol on coronary heart disease.^5^ Interventional studies on the effects of alcohol on circulating risk markers have indicated favourable effects on high-density lipoprotein (HDL) cholesterol, adiponectin and fibrinogen.^6,7^ However, accumulating evidence suggests that these markers are not causally related to cardiometabolic risk;^8-11^ the apparent cardioprotective role of modest alcohol consumption may therefore not be ascribed to mediation through these measures. Novel systemic biomarkers reflecting both alcohol intake and cardiovascular risk could serve as molecular intermediates to bridge the intricate exposure-disease relation.

The physiological effects of alcohol involve multiple metabolic pathways that extend beyond routine risk markers.^12-15^ Metabolomics technologies can provide a fine-grained snapshot of systemic metabolism and can therefore clarify the metabolic signatures of alcohol intake. Such comprehensive metabolic profiling approaches are increasingly used also in relation to alcohol research, but most biomarkers associated with alcohol intake have not been related to disease outcomes.^12-14^ Serum nuclear magnetic resonance (NMR) metabolomics enables quantitative metabolite profiling of large cohorts and biobank collections at low costs. This high-throughput methodology provides detailed lipoprotein subclass profiling, as well as quantification of fatty acids and small molecules such amino acids, of which many have recently been linked with the risk for cardiovascular disease and type 2 diabetes.^16-20^ Associations of these biomarkers with alcohol intake could therefore elucidate the beneficial and harmful metabolic processes related to alcohol consumption.

To characterize metabolic signatures of habitual alcohol consumption, we measured serum NMR metabolomics profiles from 9778 young adults from three population-based cohorts in Finland. We further assessed how change in alcohol consumption was accompanied by changes in the systemic metabolic profile. This clarifies how the detailed metabolic signature of alcohol intake tracks with changes in alcohol intake and minimizes influences of confounding. Finally, we examined the continuous shapes of the metabolic associations with alcohol intake to uncover potentially non-linear relationships with cardiovascular risk markers.

## METHODS

### Study Populations

All study participants provided written informed consent, and study protocols were approved by the local ethical committees. The study population comprised three population-based cohorts: the Northern Finland Birth Cohort of 1966 (NFBC-1966; n=4837 aged 31)^21,22^, the FINRISK 1997 study (n=2808 aged 24-45)^22,23^, and the Cardiovascular Risk in Young Finns Study (YFS; n=2133 aged 24-39 at the first time-point of metabolic profiling).^22,24^ To minimize reverse causality and to capture as far as possible physiological rather than pathological associations, analyses were restricted to individuals under 46 years of age (n=10 318 out of 13 227 indviduals with metabolic profile and alcohol data available). In addition, pregnant women (n=270), participants using lipid-lowering medication (n=17) and individuals reporting alcohol consumption >500 g/week (n=74) were excluded, yielding 9778 participants aged 24-45 for the present meta-analysis of cross-sectional associations. A subset of 1466 participants from YFS who were followed up again after 6 years (year 2001 to 2007) were assessed in longitudinal analyses.^22^ These longitudinal analyses were further validated for 1401 YFS participants with data on alcohol and metabolite levels at 10-year follow-up (year 2001 to 2011). Body mass index (BMI) and blood pressure were measured as part of the clinical examination. Current smoking status and physical activity index (assessed as metabolic-equivalent of task in NFBC-1966 and YFS, and as high or low leisure time activity in FINRISK) were assessed by questionnaires.^22^ Further details of the study populations are described in Supplementary Methods.

### Alcohol Consumption

Alcohol intake was assessed from questionnaires. In the NFBC-1966, the average volume of ethanol consumed per week was estimated from beverage-specific questions on usual weekly frequency of drinking and usual volume consumed per occasion for beer, cider and long drinks (premixed ready to drink cocktails with ethanol volume approximately 5.5%), light wine, mild, strong or home-made wine (in 16 cL glasses), and spirits (in 4 cL restaurant portions).^21^ In the FINRISK study, the volume of ethanol consumed in the past week was calculated from beverage-specific questions on consumption of beer, cider and long drinks (measured in 0.33 L bottles), wine (in 12 cL glasses) and spirits (in 4 cL restaurant portions).^25^ In the YFS, the volume of ethanol consumed in the past week was estimated from beverage-specific questions on the volume of medium beer (4.7%), strong beer (>4.7%), wine, cider/long drinks, and spirits consumed. Questions on beer, cider, and long drinks referred to 0.33 L bottles, wine to glasses and spirits to 4 cL portions.^24^ The alcohol intake was summarized as volume of ethanol in grams per week for all three cohorts, with the data harmonized to the volume of ethanol estimation across the different study questionnaires. Participants with alcohol consumption above 500 g/week (approximately 99^th^ percentile) were excluded to avoid the increased likelihood of these values being spurious due to implausible values.

### Metabolic Profiling

A high-throughput serum NMR metabolomics platform was used to quantify 76 circulating lipid and metabolite measures in the three cohorts.^26^ This metabolomics platform provides simultaneous quantification of routine lipids, lipid concentrations of 14 lipoprotein subclasses and major subfractions, and further abundant fatty acids, amino acids, ketone bodies and gluconeogenesis related metabolites in absolute concentration units (**Supplementary Table 1**). The platform has been applied extensively in epidemiological studies^16-20,22,26,27^ and details of the experimentation have been described elsewhere.^26,28^ We also analysed ten additional protein and hormonal biomarkers measured in at least two cohorts. These were high-sensitivity C-reactive protein (CRP), phospholipase activity, gamma glutamyl transferase (GGT), alanine aminotransferase (ALT), testosterone, sex-hormone binding globulin (SHBG), adiponectin, leptin, vitamin D, and insulin;^22^ they were measured using conventional clinical chemistry and mass spectrometry as described in Supplementary Methods. Inclusion of these additional measures was selected a priori because of suspected association with alcohol consumption and cardiovascular risk.

### Statistical Analysis

Metabolic measures with skewness >2 were first log_*e*_transformed. All metabolic measures were subsequently scaled to standard deviation (SD) units separately for each cohort to enable comparison of association magnitudes. Due to the correlated nature of the metabolic measures, more than 95% of the variation in the metabolomic data in all three cohorts was explained by 32 principal components. Multiple testing correction therefore accounted for 32 independent tests using the Bonferroni method, resulting in *P*<0.0016 considered statistically significant.^27^

Linear regression models were fitted for each metabolic measure with ethanol intake as the explanatory variable and the metabolite concentration as the outcome. All models were adjusted for age and sex. Results were analysed separately for each cohort and combined using fixed effect inverse-variance weighted meta-analysis after confirming the consistency across the three cohorts. Sensitivity analyses were conducted with additional adjustment for BMI, smoking, and physical activity. Association magnitudes are presented combined for men and women in the main text since association magnitudes were generally similar for both sexes; sex-stratified analyses are presented in the Data Supplement.

Data on alcohol consumption and metabolite levels were available for 1466 individuals from YFS at 6-year follow-up; this was used to examine whether changes in metabolite concentrations track with changes in alcohol consumption during follow-up. For these longitudinal analyses, linear regression models were fitted with 6-year change in metabolite concentration as the outcome and 6-year change in ethanol intake as the explanatory variable with adjustment for age and sex. Baseline and follow-up metabolite concentrations were standardized to SD-units at baseline, so that cross-sectional and longitudinal associations are directly comparable. The resemblance between the overall cross-sectional and longitudinal association patterns was quantified using the *R*^2^ goodness-of-fit and the slope of the linear fit.^1,22^ To further validate the consistency between the cross-sectional and longitudinal associations, the association patterns were also compared using *R*^2^ for 1401 individuals with alcohol and metabolite data at 10-year follow-up.

The continuous shape of the metabolic associations with alcohol consumption were examined descriptively using local quadratic regression fitting, with each smoothing function segment evaluated at 25 points through the range of ethanol intake. To adjust for age and sex, the absolute concentration of each metabolic measure was first regressed for age and sex in each cohort and the resulting residuals were pooled prior to fitting and rescaled to absolute units. Equivalent analyses were done stratified by sex. Statistical analyses were conducted using R 3.1.

## RESULTS

The study population comprised 9778 young adults from three Finnish population-based cohorts with comprehensive lipid and metabolite profiling data. Clinical characteristics of the three cohorts are shown in **Table 1**. The mean age was 33 years (range 24–45) and 53% of the participants were women. The median consumption of alcohol was around 3 drinks per week (36 grams of ethanol) in all three cohorts. Despite differences in the questionnaires on alcohol usage relating to usual intake or consumption during the week prior to the clinical examination, the distribution of alcohol intake was similar across the three cohorts (**Supplementary Figure S1**). Mean concentrations of the metabolic measures are listed in **Supplementary Table 1**.

**Table 1.** Clinical characteristics of the study population.

| Characteristic | Northern Finland Birth Cohort of 1966 | FINRISK 1997 study | Cardiovascular Risk in Young Finns Study |
| --- | --- | --- | --- |
| Number of participants (men/women) | 4837<br>(2414/2423) | 2808<br>(1320/1488) | 2133<br>(973/1160) |
| Age [yr] | 31.2 (0.4) | 35.6 (6.2) | 31.7 (5.0) |
| Body mass index [kg/m<sup>2</sup>] | 24.6 (4.1) | 25.2 (4.3) | 25 (4.4) |
| Systolic blood pressure [mmHg] | 125 (13) | 127 (15) | 117 (13) |
| Total cholesterol [mmol/L] | 5.1 (1.0) | 5.1 (1.0) | 5.1 (1.0) |
| HDL cholesterol [mmol/L] | 1.5 (0.4) | 1.4 (0.3) | 1.3 (0.3) |
| Triglycerides [mmol/L] | 1.0 [0.8-1.4] | 1.0 [0.7-1.4] | 1.0 [0.8-1.5] |
| Plasma glucose [mmol/L] | 5 [4.7-5.3] | 5.0 [4.7-5.3] | 4.8 [4.6-5.2] |
| Insulin [IU/L] | 6.7 [4.9-8.9] | 7.5 [6.2-9.4] | 4.7 [3.3-6.7] |
| Smoking prevalence [%] | 42% | 30% | 24% |
| Alcohol usage [g/week] | 32.6 [11.5-76.1]<sup>a</sup> | 36.7 [0-93.6]<sup>b</sup> | 39 [0-103]<sup>b</sup> |
| Alcohol usage [drinks/week] | 2.6 [0.9-6.1]<sup>a</sup> | 3.0 [0.0-7.6]<sup>b</sup> | 3.1 [0.0-8.3]<sup>b</sup> |

Values are mean (SD) and median [interquartile range] for normally distributed and skewed variables, respectively.
<sup>a</sup> Based on usual volume of ethanol consumed.
<sup>b</sup> Based on volume of ethanol consumed in the past week.

### Alcohol associations with lipoprotein lipids

Cross-sectional associations of alcohol consumption with 38 lipoprotein and lipid measures are shown in the left panel of **Figure 1**; the corresponding associations of change in alcohol consumption with change in lipid concentrations are shown in the right panel. The overall cross-sectional association pattern was recapitulated in the longitudinal association pattern: the lipids most strongly associated with alcohol consumption also displayed the highest responsiveness to changes in alcohol intake during follow-up. Increased alcohol consumption was modestly associated with elevated lipid levels within the largest very-low-density lipoprotein (VLDL) subclasses, whereas associations with lipids in the low-density lipoprotein (LDL) subclasses appeared very weak. However, increased alcohol intake was robustly associated with larger VLDL particle size, while LDL particle size displayed a strong inverse association. Prominent associations were observed between higher alcohol consumption and higher lipid concentrations for all high-density lipoprotein (HDL) subclasses, particularly for the medium-sized and small HDL particles. Concomitantly, the cholesterol and phospholipid concentrations in the HDL subclasses were robustly elevated in relation to higher alcohol intake, whereas triglyceride in HDL were only modestly elevated. When examining the continuous association shapes, HDL-related measures were broadly linear or modestly declining in slope across the range of alcohol consumption, while more complex continuous association shapes were observed for the apolipoprotein B-carrying lipids (**Supplementary Figure 2**). Here, inverse associations were observed in the first segment up to 50 g/week, with convex association curves for men and declining slopes for women. The non-linear association shapes for these lipid measures partly mask the associations in the linear models. The pattern of lipid associations with alcohol intake was highly coherent across all three cohorts analysed (**Supplementary Figure S3**). Association magnitudes in absolute concentration units are listed in **Supplementary Table S1**. For example, 100 g higher ethanol intake per week (corresponding to about 8 drink units; 1.29 SD) was associated with 0.073 mmol/L higher HDL cholesterol.

**Figure 1:**
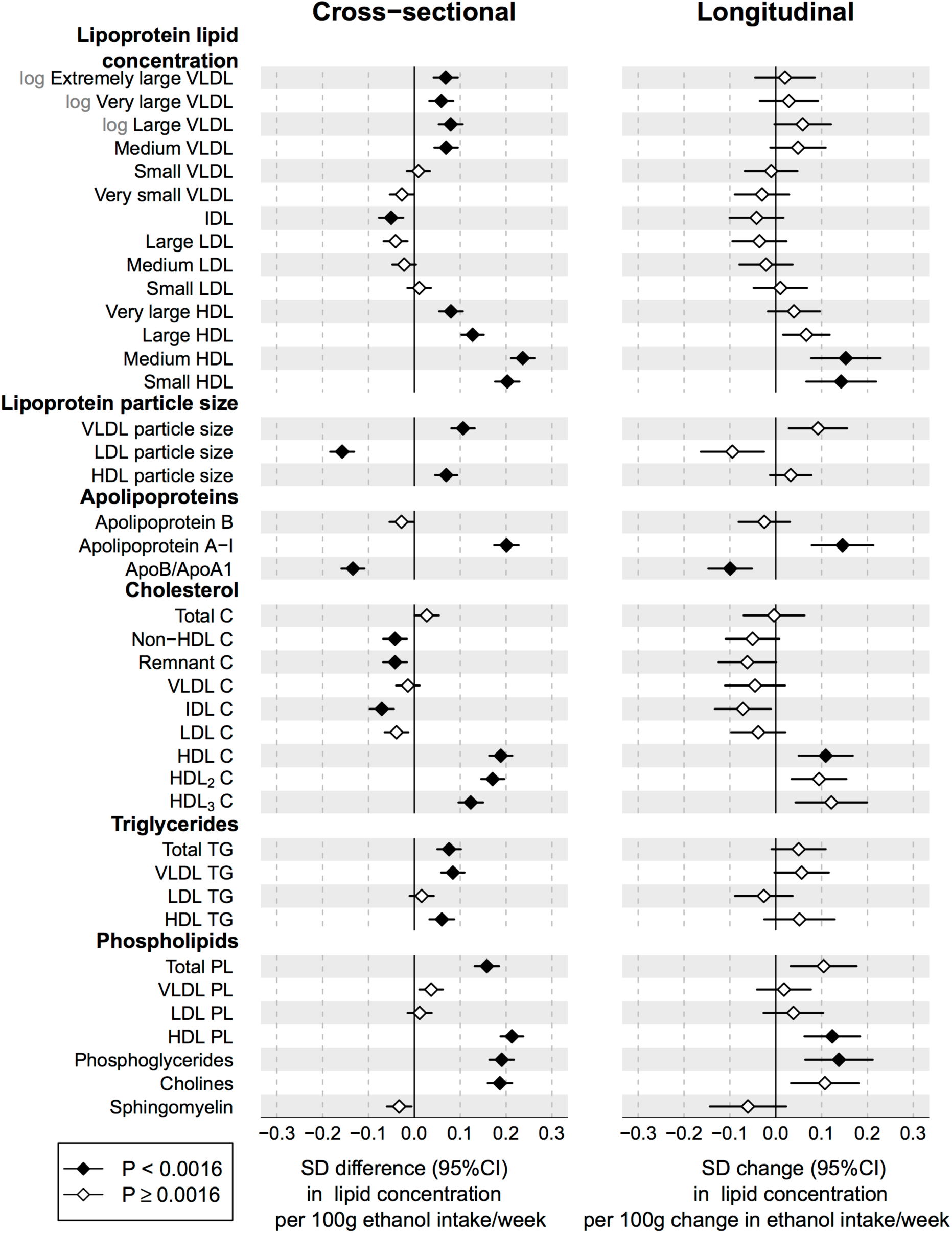
Cross-sectional and longitudinal associations between alcohol consumption and lipoprotein lipid measures. *Left*: Cross-sectional associations of alcohol intake with lipid measures m eta-analyzed for three cohorts of young adults (n=9778). *Right*: Changes in lipid concentrations associated with change in alcohol intake after 6-year follow-up for 1466 individuals. All associations were adjusted for age and sex. Error bars denote 95% confidence intervals. Association magnitudes in absolute concentration units are listed in **Table S1**. Continuous shapes of the metabolic associations with alcohol intake are shown in **Supplementary Figure S2**.

### Alcohol associations with fatty acids

The cross-sectional and longitudinal associations of alcohol consumption with circulating fatty acid levels are shown in **Figure 2**. The association pattern observed in the cross-sectional analyses resembled that seen within person over the 6-year follow-up period, except for omega-3 fatty acid related measures. Higher alcohol intake was robustly associated with an increase in the absolute concentrations of total fatty acids, saturated fatty acids and monounsaturated fatty acids (MUFA). In contrast, absolute concentrations of omega-6 fatty acids were not associated with alcohol consumption. However, the proportions of fatty acid levels to total fatty acids are often considered better indicators of metabolic risk.^1,2,16^ These measures displayed a pattern of strongly elevated MUFA ratio with higher alcohol consumption, whereas omega-6 fatty acid ratio was strongly inversely associated with alcohol intake. These fatty acid associations were linear across the range of alcohol intake. The association magnitudes of MUFA ratio (0.26 SD higher per 100 g weekly ethanol intake, equivalent to 0.90%) and omega-6 proportion (-0.23 SD lower per 100 g weekly ethanol intake, equivalent to 0.87%) were stronger than that of HDL cholesterol. The association of alcohol intake with the proportion of omega-3 fatty acid was only weakly positive cross-sectionally and inconsistent with this longitudinally.

**Figure 2:**
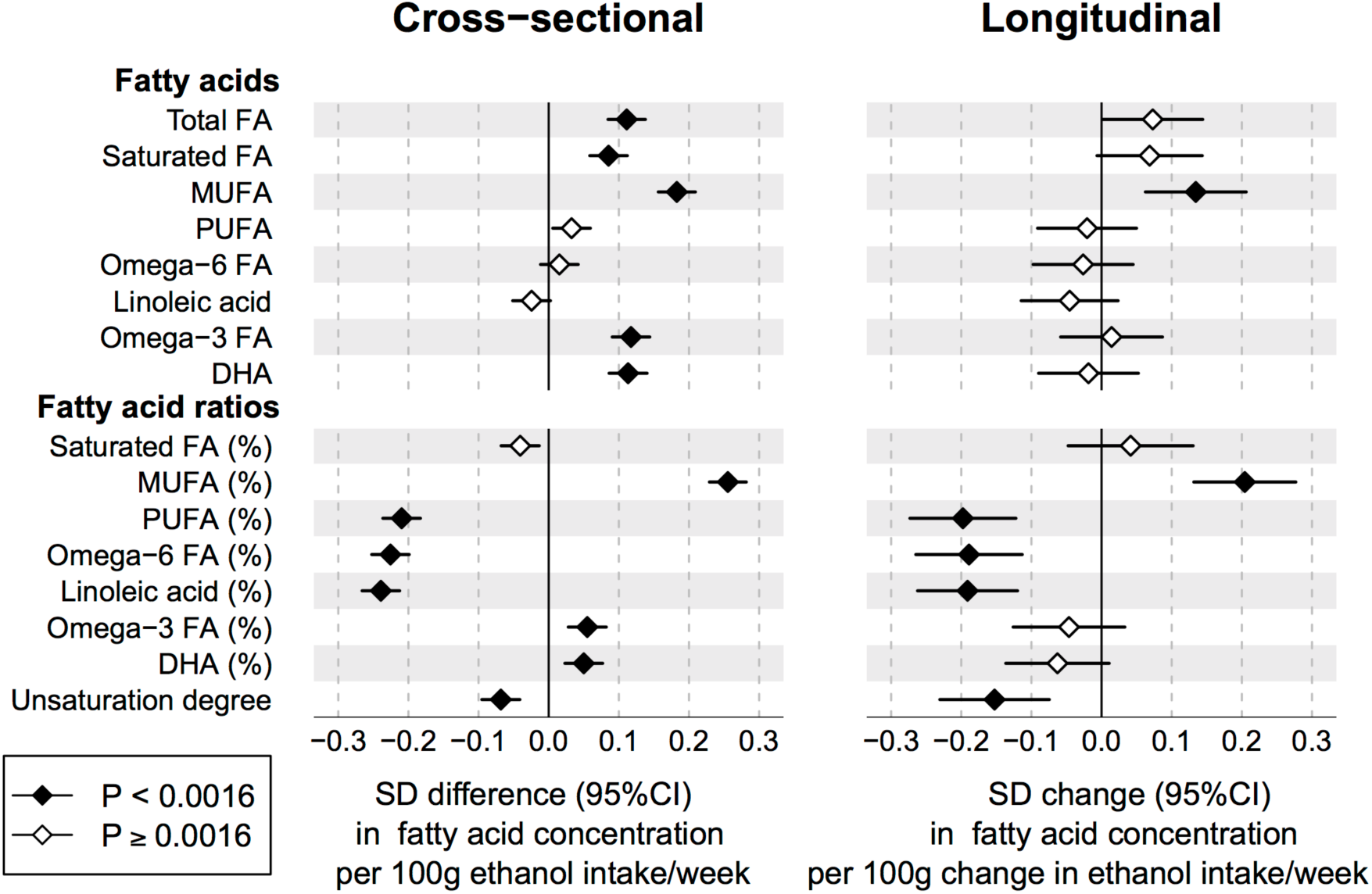
Cross-sectional and longitudinal associations between alcohol consumption and fatty acids. *Left*: Cross-sectional associations of alcohol intake with circulating fatty acids meta-analyzed for three cohorts of young adults (n=9778). *Right*: Changes in fatty acid levels associated with change in alcohol intake after 6-year follow-up for 1466 individuals. All associations were adjusted for age and sex. Error bars denote 95% confidence intervals.

### Alcohol associations with metabolites and hormones

The cross-sectional and longitudinal associations of alcohol consumption with circulating amino acids, gluconeogenesis metabolites, and various inflammatory and hormonal measures are show in **Figure 3**. The strongest associations for low-molecular-weight metabolites were observed for glutamine and citrate, both inversely associated with increased alcohol intake. While most of the small molecule metabolites were not strongly associated with alcohol consumption based on the linear models, subtle nonlinear association shapes were evident for several measures, e.g. phenylalanine and glycerol. Higher alcohol intake was modestly associated with increased levels of several amino acids, glycolysis and gluconeogenesis related metabolites as well as glycoprotein acetyls and C-reactive protein, markers of chronic inflammation. Higher testosterone was associated with alcohol consumption for both men and women, while an inverse association with SHBG was only observed for men. Higher alcohol intake was also associated with the adipokine adiponectin and elevated circulating levels of the liver enzymes GGT and ALT. Although the power to denote statistical significance was limited, the changes in metabolite and hormonal concentrations associated with 6-year change in alcohol consumption broadly matched with the cross-sectional associations.

**Figure 3:**
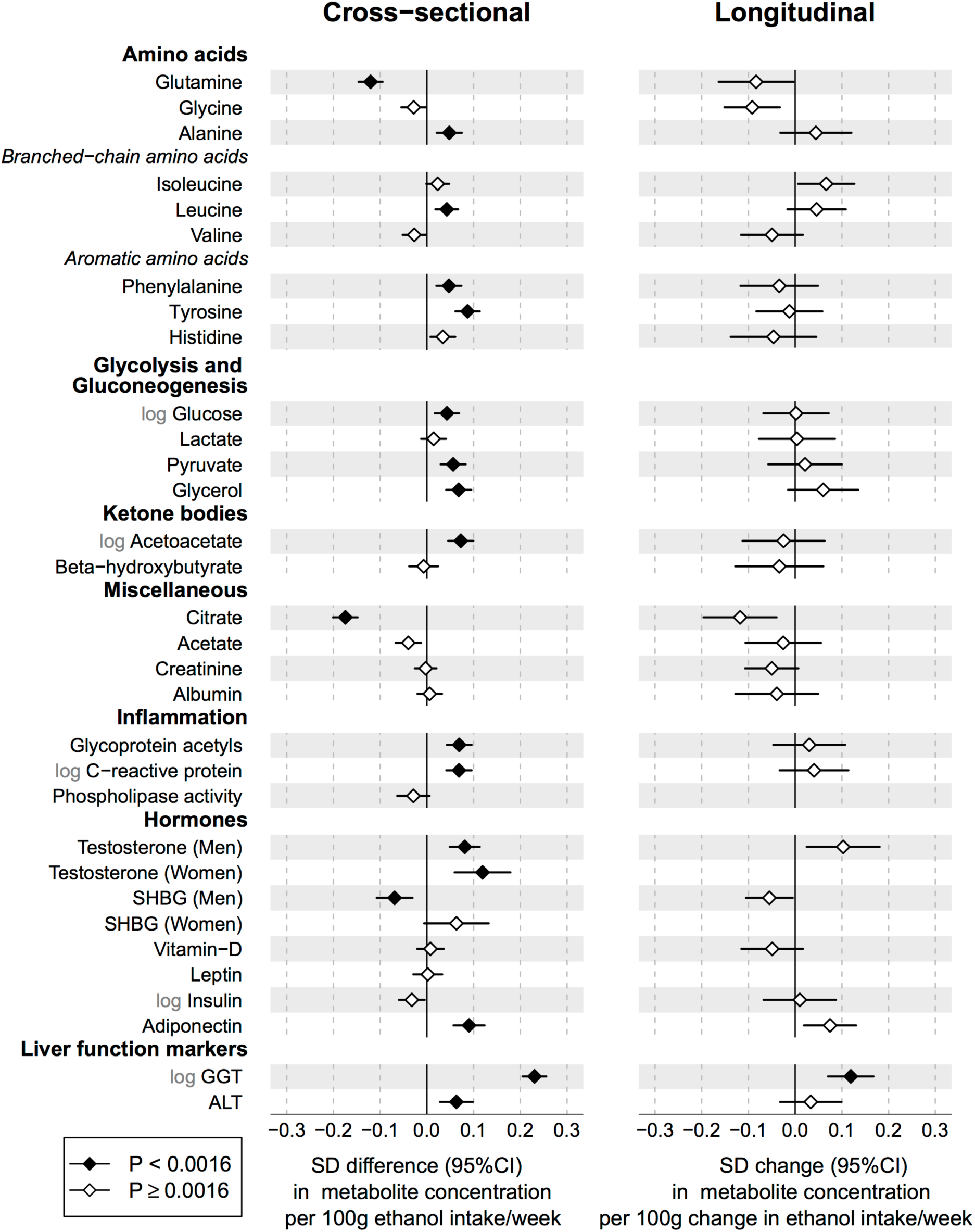
Cross-sectional and longitudinal associations of alcohol consumption with low-molecular-weight metabolites and hormonal measures. *Left*: Cross-sectional associations of alcohol intake with metabolite and hormonal measures meta-analyzed for three cohorts of young adults (n=9778). *Right*: Changes in metabolite and hormone levels associated with change in alcohol intake after 6-year follow-up for 1466 individuals. All associations were adjusted for age and sex. Error bars denote 95% confidence intervals.

### Consistency of cross-sectional and longitudinal association patterns

The resemblance between the overall metabolic association patterns of alcohol consumption observed cross-sectionally and longitudinally is illustrated in **Figure 4**. The association magnitudes followed a straight line, with goodness of fit *R*^2^=0.83 (95% confidence intervals 0.77-0.90). The slope of the linear fit was 0.72±0.04, indicating that each unit change in alcohol intake was on average accompanied by a slightly weaker change in the metabolic profile compared to that estimated from the cross-sectional analyses. The consistency between the cross-sectional and longitudinal associations was even stronger if only comparing measures associated with alcohol intake at *P*<0.0016 (*R*^2^=0.87). The match of the association patterns was also similar when cross-sectional and longitudinal associations were compared only for the same 1466 individuals (*R*^2^=0.79). Data on metabolic profiling and alcohol intake were also available at 10-year follow-up from the same study population;^3-5,22^ highly coherent results were obtained when the longitudinal and cross-sectional association patterns were compared at this extended follow-up duration (*R*^2^=0.77).

**Figure 4:**
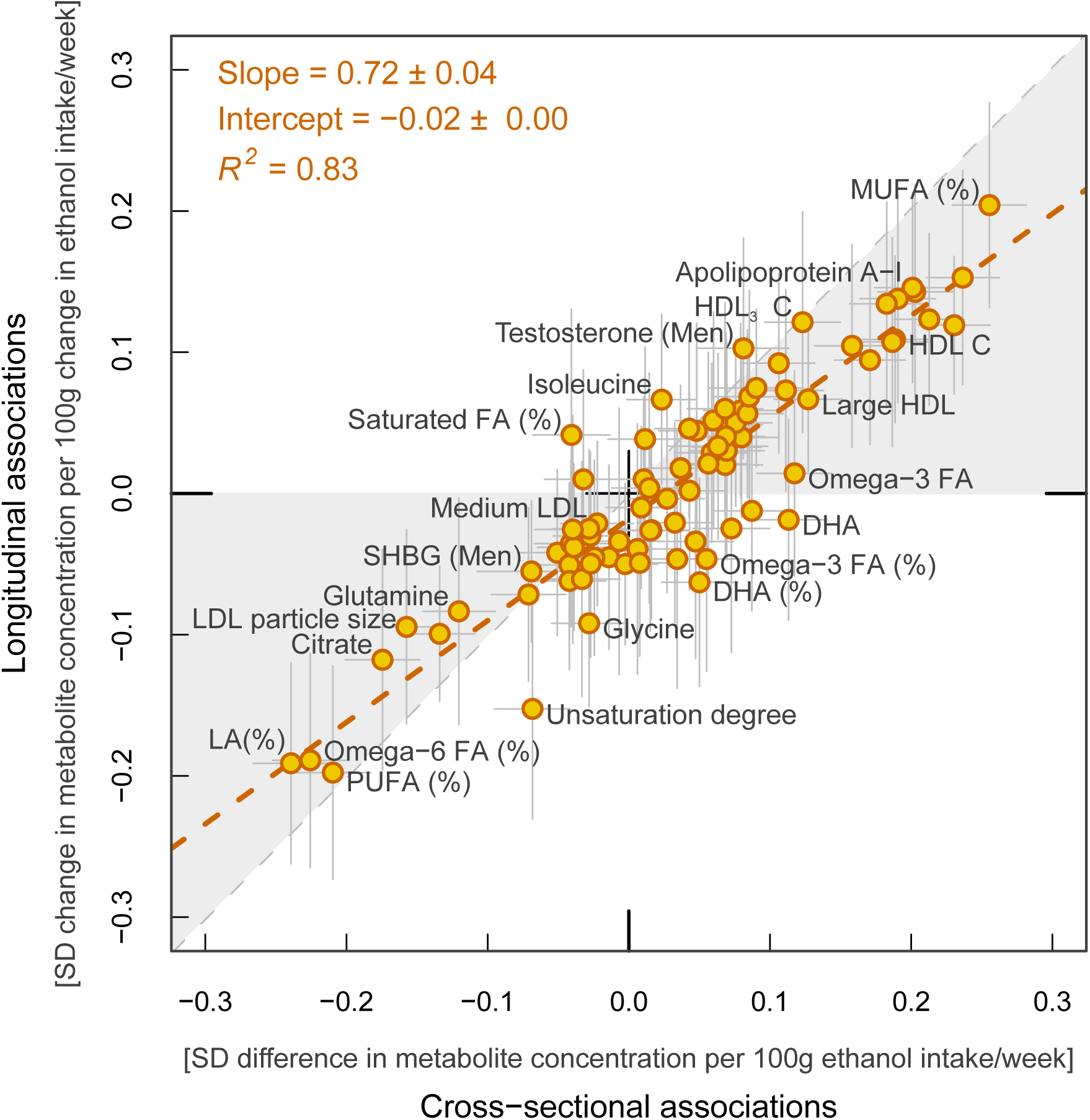
Consistency between metabolic associations with alcohol intake at a single time-point with the corresponding metabolic changes associated with 6-year change in alcohol consumption. The resemblance between the overall association patterns was quantified by the linear fit between the cross-sectional and longitudinal associations with alcohol intake (red dashed line).

### Sensitivity analyses

The cross-sectional associations of alcohol intake with metabolic measures were generally concordant across the three cohorts (Supplementary Figure S3). Men and women displayed broadly similar metabolic association patterns with alcohol intake (**Supplementary Figure S4**). Association magnitudes on the absolute scale tended to be stronger for women, but if taking the higher variation in alcohol intake for men into account then association magnitudes were on average 12% stronger for men. Although some sex differences were apparent from the linear modelling, e.g. inverse associations for LDL subclasses for women only, these were mostly related to subtle differences in the continuous association shapes. Surprisingly, adjustment for BMI, smoking status, and physical activity index had little affect on strength of association (3% attenuation on average; **Supplementary Figure S5**). The results were also similar if additional adjustment for baseline alcohol was made in the longitudinal modelling. All results were essentially unaltered if excluding the 1388 individuals who abstained from alcohol consumption according to the questionnaires.

### Continuous shape of metabolic associations with alcohol

The shapes of the metabolic associations as a function of alcohol consumption are shown for all 86 metabolic measures in Supplementary Figure S2. The continuous association shapes for 12 selected measures are illustrated in **Figure 5**. For most of the metabolic measures strongly correlated with alcohol intake, the shapes of association were approximately linear. This is exemplified for MUFA and omega-6 ratios, relative to total fatty acids, as well as glutamine. Other metabolic measures displayed non-linear association shapes, with strong initial increase (e.g., medium HDL lipids) or decrease (e.g., citrate) that became less steep at high ranges of alcohol consumption. Omega-3 ratio displayed a steep increase followed by a plateau association. A substantial number of metabolic measures displayed more complex nonlinear associations, as exemplified for medium VLDL and medium LDL lipid concentrations in Figure 5. Total triglycerides and LDL cholesterol, respectively, displayed similar shapes as these subclass measures. U-shaped curves for modest alcohol intake (up to 200g ethanol/week, approximately 2 drinks per day; 93% of the study population in this range) were particularly prominent for men in the case of atherogenic lipid measures (Supplementary Figure S2). However, similar convex shapes were observed for both men and women in this range of alcohol intake for other cardiometabolic risk biomarkers, such as phenylalanine.

**Figure 5:**
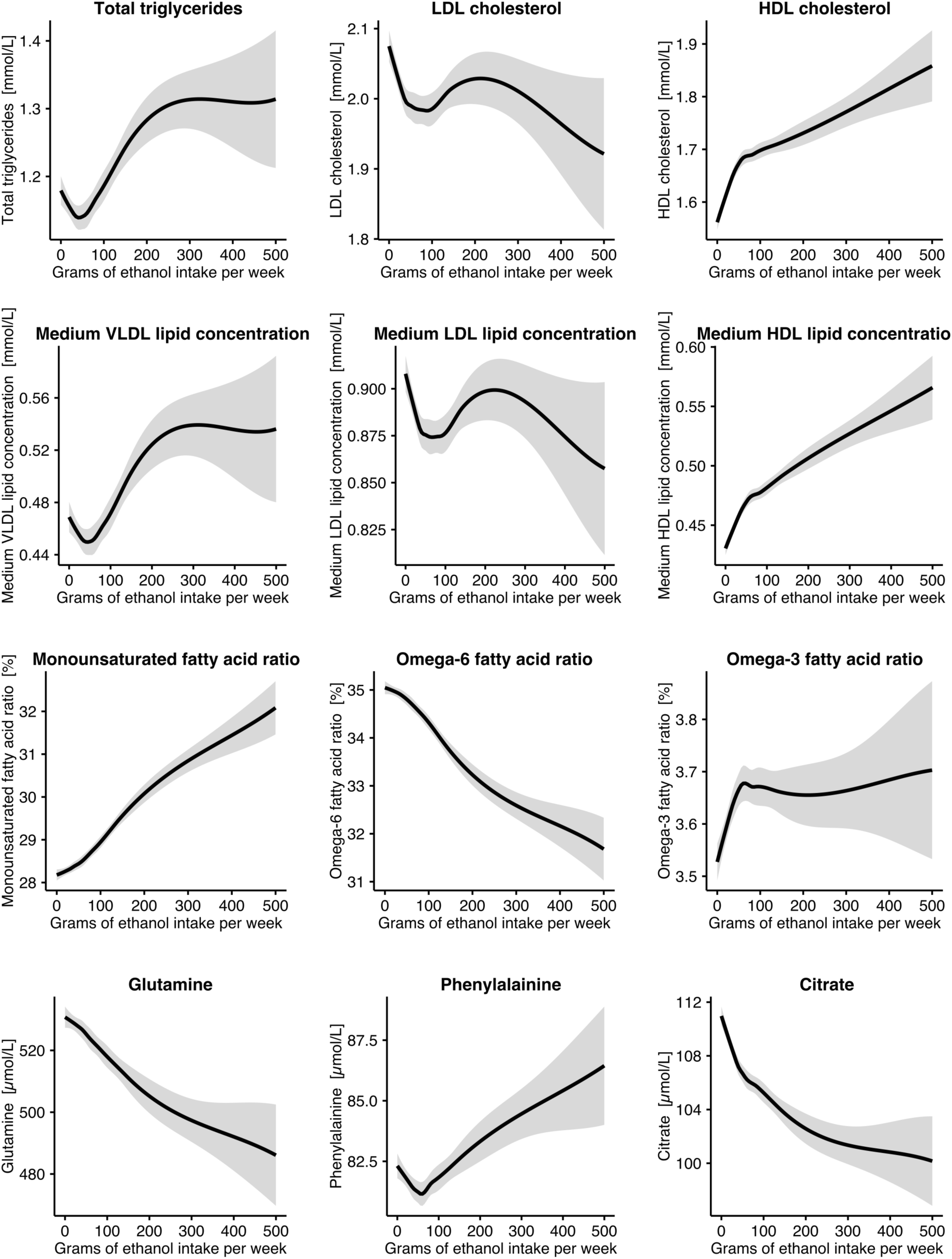
Lipid and metabolite concentrations as a function of alcohol consumption for 12 selected metabolic measures (n=9978). The shaded curves denote the 95% confidence intervals of the local polynomial regression fits. Associations were adjusted for age, sex and cohort. The continuous shapes of associations for all 86 metabolic measures are shown for men and women in **Supplementary Figure S2**.

## DISCUSSION

Metabolic profiling of almost 10,000 young adults revealed an intricate pattern of lipid and metabolite associations with usual alcohol consumption. The molecular signature linked with alcohol intake cover multiple metabolic pathways, including lipoprotein subclasses, fatty acid composition, glutamine and citrate regulation, and hormonal balance. These metabolic perturbations comprise a mixture of both favourable and adverse effects in relation to cardiovascular disease and all-cause mortality risk.^3,16,17^ The pattern of metabolic changes accompanying change in alcohol intake at 6-year follow-up was highly consistent with the metabolic association pattern observed at a single time-point, suggesting that the metabolomic signature of alcohol consumption tracks with long-term changes in alcohol use. Although potential causal disease relations remain unclear for many of the metabolic biomarkers, our findings provide improved understanding of the diverse molecular processes related to alcohol intake, and may help to clarify the complex mediation between alcohol and cardiovascular risk.

The molecular underpinnings of the apparent cardioprotective effects of modest alcohol consumption reported in observational studies remains contentious.^3,5,6,24^ Numerous observational and interventional studies have found increased HDL cholesterol and apolipoprotein A-I levels with higher alcohol intake.^3,6,7,29^ These lipoprotein measures serve here as positive controls, along with the liver enzymes markers, and set the association magnitudes into perspective: several of the novel metabolic biomarkers were as strongly associated with alcohol intake as HDL cholesterol and GGT. Nonetheless, accumulating evidence argue against a causal role of HDL cholesterol in cardiovascular disease,^8-11^ so other underpinning mediators may be required to explain potential cardioprotective effects of moderate alcohol consumption. Medium-sized and small HDL subclasses displayed the most pronounced associations, whereas the association slopes for larger HDL subclasses levelled off after initial rapid increase. However, the lipid levels of medium and small HDL are weaker biomarkers of lower cardiovascular risk than that of large HDL.^12-16,30^ If changes in HDL metabolism due to alcohol intake are reflecting some underlying cardioprotective mechanism, then the effects appear less favourable than assumed by assessing only HDL cholesterol since this measure combines subclasses with heterogeneous associations.

In contrast to HDL cholesterol, the causal atherogenic role of LDL particles is well established and increasingly clear also for the triglyceride-rich lipoprotein particles.^12-14,31^ These apolipoprotein B-carrying particles displayed weak or no association with alcohol intake in the standard linear models; however, inverse associations were observed for women in sex-stratified analyses and men displayed U-shaped associations, with most favourable lipid levels observed around 50 grams of ethanol intake per week. Similar sex differences have been observed previously for LDL cholesterol.^16-20,32,33^ Although only a small fraction of the study population were heavy drinkers, we recapitulated the anticipated low levels of LDL subclass lipids at the highest alcohol range. However, LDL particle size strongly decreased along with increasing alcohol intake, and some studies have related this measure to higher cardiovascular risk.^21,22,30^ These results illustrate how lipoprotein subclass profiling may uncover potentially unfavourable risk effects missed by routine biomarkers. Further, the nonlinear shapes underline the need to assess continuous associations with alcohol intake for both established and emerging biomarkers of cardiometabolic risk. The more favourable circulating levels of atherogenic lipids observed for individuals with modest alcohol consumption may partly contribute to explain the apparent cardioprotective effect for moderate drinkers; however, the association magnitudes were too weak to fully attribute the consistent reports of the U-shaped association with cardiovascular risk to these lipid measures.

The relation between alcohol intake and the fatty acid balance revealed a mixed set of associations, with mostly adverse aberrations in terms of biomarkers for cardiovascular risk.^16,22,23^ Subtle sex differences and non-linear association shapes contributed to the complex picture, e.g. for omega-3 fatty acid ratio. MUFA ratio displayed the strongest association with alcohol intake among all the 86 metabolic measures analysed. Contrary to the dietary intake of this measure, higher circulating MUFA ratio is a biomarker of higher cardiovascular and diabetes risk.^16,22,24,18,22^ Likewise, the robust association of alcohol intake with lower proportion of omega-6 fatty acids is also related to higher cardiometabolic risk.^16,18,22,34^ Our findings are consistent with much smaller studies on men at high-risk for cardiovascular disease or suffering from alcohol dependence.^21,35,36^ The prominent fatty acid associations were essentially linear across the entire range of alcohol consumption and were corroborated by the longitudinal analyses. Although the causality of these fatty acids remains unclear, the primarily adverse changes in these emerging risk markers indicate also unfavourable metabolic aberrations with modest alcohol consumption.

Recent metabolic profiling studies have linked several circulating amino acids and other small molecules with the risk for cardiovascular disease, type 2 diabetes and all-cause mortality.^16,17,19,25,37,38^ Dietary determinants of these circulating biomarkers remain poorly understood. In the present analyses, only few low-molecular-weight metabolites were strongly associated with alcohol intake. The most prominent associations were for citrate and glutamine, both strongly inversely associated with alcohol intake but otherwise weakly correlated with established risk factors.^17,22,24^ Our results are supported by a smaller metabolomics study of alcohol intake in Japanese men.^14,26^ Higher citrate levels have been linked with modestly lower risk for cardiovascular disease,^16-20,22,26,27^ but simultaneously with higher risk of all-cause mortality.^17,26,28^ The biological underpinnings of citrate’s links with disease remain elusive. Higher circulating glutamine is associated with lower risk for cardiovascular disease^16,22^ and type 2 diabetes.^19,27,37^ Glutamine and citrate are connected via the citric acid cycle and both metabolites play critical roles at the centre of cancer cell metabolism.^39,40^ Multiple other metabolites and hormones also displayed associations with alcohol intake; however, further investigations are required to clarify whether these molecular perturbations could arise as a combined effect of alcohol intake on multiple metabolic pathways.

Our study has both strengths and limitations. The large sample size provides robust evidence for many novel metabolic markers of alcohol consumption, in particular since results were coherent across the three cohorts despite differences between the measures of alcohol intake. Although our study was conducted solely in the homogenous Finnish population, many of the strongest metabolic biomarkers have been related to alcohol intake in smaller cross-sectional studies of populations with other ethnicities.^12,14,36^ Observational studies on alcohol consumption are susceptible to confounding and reverse causation.^3,5^ By design, all study participants were young and relatively healthy, which minimizes influences of reverse causality. The associations were also essentially unaltered when excluding non-drinkers; the results are thus unlikely to be influenced by those who may have stopped drinking for health reasons. All results were similar when adjusting for adiposity, smoking and physical activity, suggesting limited confounding by these risk factors clustering with alcohol intake. The lack of association with leptin, and the distinct metabolic association pattern from that caused by elevated BMI^22^, make it implausible that the metabolic perturbations would be mediated via effects of alcohol on adiposity. However, dietary composition may potentially confound the results. The consistency of the cross-sectional and longitudinal association patterns helps exclude influence of unmeasured confounders that are fixed individual characteristics; however, we acknowledge that also the longitudinal associations remain susceptible to other types of confounding. The majority of study participants consumed alcohol only in low or moderate amounts, which prevents inferences about the metabolic associations with heavy alcohol consumption. Metabolomics of interventional studies could clarify whether the diverse metabolic changes arise due to acute effects of alcohol intake or chronic exposure, for instance mediated via impaired liver function. Further studies may also characterize the metabolic effects of binge drinking, specific beverage types, and whether the circulating markers of alcohol consumption could benefit identification of individuals at risk for alcoholic liver disease.

In conclusion, metabolic profiling of three large population-based cohorts identified novel metabolic markers for dose-dependent alcohol intake. The detailed metabolic phenotyping further clarified the association shapes for numerous established biomarkers related to alcohol and cardiometabolic risk. The metabolic signature of alcohol consumption included molecular perturbations linked with both higher and lower cardiovascular risk. Many metabolic measures displayed an optimum level at modest alcohol intake. The striking match between the overall cross-sectional and longitudinal association patterns provides evidence in support of the metabolic changes arising, at least partly, due to alcohol consumption. Comprehensive metabolic profiling in these large cohorts thus elucidated the metabolic influences of alcohol consumption and clarified the double-edged relation between alcohol and cardiometabolic biomarkers.

## DISCLOSURES

PW, AJK, PS, and MAK are shareholders of Brainshake Ltd, a company offering NMR-based metabolite profiling. PW, QW, MT, TT, AJK, and PS report employment relation for Brainshake Ltd. No other authors reported disclosures.

## SOURCES OF FUNDING

This study was supported by Strategic Research Funding from the University of Oulu, Sigrid Juselius Foundation, Academy of Finland, Novo Nordisk Foundation, Paavo Nurmi Foundation, Yrjö Jahnsson Foundation, Emil Aaltonen Foundation, Finnish Diabetes Research Foundation, Finnish Foundation for Cardiovascular Research, and UK Medical Research Council via the University of Bristol Integrative Epidemiology Unit (MC_UU_12013/1 and MC_UU_12013/5). The Cardiovascular Risk in Young Finns Study is supported by Academy of Finland, the Social Insurance Institution of Finland; Kuopio, Tampere and Turku University Hospital Medical Funds, Juho Vainio Foundation, Paavo Nurmi Foundation, Finnish Foundation of Cardiovascular Research, Finnish Cultural Foundation, Emil Aaltonen Foundation, and Yrjö Jahnsson Foundation. The Northern Finland Birth Cohort has received financial support from Academy of Finland, University Hospital Oulu, Biocenter Oulu, University of Oulu, the European Commission (EURO-BLCS, Framework 5 award QLG1-CT-2000-01643, ENGAGE project and grant agreement HEALTH-F4-2007-201413, EurHEALTHAgeing (277849), European Regional Developmental Fund), EU H2020-PHC-2014 (Grant no. 633595), NHLBI grant 5R01HL087679-02 through the STAMPEED program (1RL1MH083268-01), NIH/NIMH (5R01MH63706:02), Stanley Foundation, UK Medical Research Council, and Wellcome Trust.

